# NMDA-receptor-Fc-fusion constructs neutralize anti-NMDA receptor antibodies

**DOI:** 10.1101/2022.04.29.490085

**Authors:** Stephan Steinke, Toni Kirmann, Jana Nerlich, Iron Weichard, Philip Kuhn, Torsten Bullmann, Andreas Ritzau-Jost, Filiz Sila Rizalar, Harald Prüss, Volker Haucke, Christian Geis, Michael Hust, Stefan Hallermann

**Author notes:** These authors contributed equally (Stephan Steinke, Toni Kirmann).

## Abstract

N-methyl-D-aspartate receptor (NMDAR) encephalitis is the most common subtype of autoimmune encephalitis characterized by a complex neuropsychiatric syndrome ranging from memory impairment and psychosis to coma. Patients develop an intrathecal immune response against NMDARs with antibodies that presumably bind to the amino-terminal domain (ATD) of the GluN1 subunit. The therapeutic response to immunotherapy is often delayed and does not directly interfere with intrathecal synthesis of pathogenic antibodies. Therefore, new therapeutic approaches for fast neutralization of NMDAR antibodies are needed. Here, we developed fusion constructs consisting of the Fc part of immunoglobulin G and the ATDs of either GluN1 or GluN2B or both, GluN1 and GluN2B, subunits. Surprisingly, both subunits were required to generate high-affinity epitopes. The construct with both subunits efficiently prevented NMDAR binding of patient-derived monoclonal antibodies and of patient cerebrospinal fluid containing high-titer NMDAR antibodies. Furthermore, it inhibited the internalization of NMDARs in rodent dissociated neurons and human induced pluripotent stem cells (iPSC)-derived neurons. Finally, the construct stabilized NMDAR currents recorded in rodent neurons. Our results demonstrate that both GluN1 and GluN2B subunits contribute to the main immunogenic region of the NMDAR and provide a promising strategy for fast and specific treatment of NMDAR encephalitis, which can complement immunotherapy.

## Introduction

Since the initial description of anti-NMDAR encephalitis in 2007 (ref.^1^), the impact in neurology and psychiatry has been remarkable and led to the definition of a new group of CNS disorders called ‘autoimmune encephalitis’. These diseases are characterized by specific and pathogenic antineuronal antibodies directed at synaptic antigens.^2^ Patients with autoimmune encephalitis present with a variable pattern of severe neuropsychiatric symptoms that may lead to long-lasting coma within weeks.^3^ Potential triggers for the autoimmune response include paraneoplastic molecular mimicry by ectopic expression of neuronal antigens in tumors, e.g. teratomata in NMDAR encephalitis, or antigens released by neuronal damage, e.g. after herpes simplex encephalitis. However, in the majority of cases no such triggers have been identified yet. The immune response in NMDAR encephalitis induces circulating B cells and intrathecally expanded antibody-producing cells in the brain.^2,4^ The main pathomechanism is binding of the antibodies to the amino terminal domain (ATD) of the NMDAR GluN1 subunit ^5,6^, which causes clustering and internalization of NMDARs, most likely by cross-linking mechanisms.^7-9^

For the treatment of NMDAR encephalitis, no validated guidelines exist yet and the available immunotherapy shows limited efficacy (reviewed by Sell et al.^10^). About 25% of patients with NMDAR encephalitis are refractory to treatment^11^ and often require long-term intensive care treatment due to life-threatening complications.^12^ Antibody-depleting strategies (e.g. plasma exchange), B cell depletion (e.g. with rituximab), or experimental plasma cell targeting (e.g. with the proteasome inhibitors bortezomib^13,14^) have limited efficiency in the CNS compartment^15^ and/or do not affect the main antibody-producing intrathecal plasma cells directly.^3^ Furthermore, due to the half-life of immunoglobulin G (IgG) antibodies of ∼20 days and long-living plasma cells in the CNS compartment, the therapeutic response to any immunotherapy is significantly delayed. These limitations of immunotherapy explain the often prolonged recovery from disease symptoms and indicate that more specific and effective therapeutic approaches are needed.

The following three more specific approaches to interfere with direct effects of anti-NMDAR antibodies have been considered. First, the activation of the ephrinB2 receptor (EphB2R) by a soluble form of its ligand ephrin-B2 is able to stabilize NMDAR density^7^ and to rescue visuospatial learning^16^ after pathogenic anti-NMDAR antibody application in mice. The underlying mechanism involves phosphorylation of the GluN2B subunit of the NMDAR upon activation of the EphB2R^17^, which controls synaptic NMDAR clustering and retention.^18^ Secondly, positive allosteric modulators of NMDAR, such as 24(S)-hydroxycholesterol,^19^ were shown to potentiate the remaining, non-internalized NMDAR, thereby compensating for the NMDAR loss.^20,21^ However, both approaches might overcompensate the loss of NMDARs and interfere with endogenous NMDAR function and membrane cycling. Thirdly, monovalent Fab fragments were able to bind to NMDARs without inducing cross-linking, internalization, or reduction of NMDAR density.^9,22^ However, interference with Fab fragments to prevent binding of the antibodies is hampered by the greater avidity of IgG in comparison to Fab fragments, a potential pathogenic effect of Fab fragments by interfering with NMDAR function^22^, and a shorter serum half-life of Fab fragment due to the protective function of the lacking Fc fragment.^23^ Thus, the currently considered approaches to specifically treat NMDAR encephalitis have certain limitations.

To overcome these limitations, we developed an ATD-Fc-fusion construct that can neutralize pathogenic autoantibodies of patients while leaving NMDAR function unperturbed. We show that the construct prevents the binding of autoantibodies to the NMDAR, which represents the initial, disease-defining step. The constructs can therefore inhibit the hallmarks of the disease’s pathophysiology including internalization of NMDARs and reduction of NMDAR currents.

## RESULTS

### An engineered ATD-Fc-fusion construct has high affinity for pathogenic anti-NMDAR antibodies

The production of stable epitopes for pathogenic anti-NMDAR autoantibodies is complicated by the hydrophobic nature of the NMDAR and the conformation-dependent epitope in the ATD of NMDARs.^5,24^ We therefore designed several fusion constructs containing different parts of the ATD of NMDARs and a Fc fragment of mouse or human IgG (Fig. 1A). The ATD part contained either two GluN1 ATDs from amino acid (aa) 19 to aa 397, two GluN2B ATD from aa 25 to aa 395, or a combination of two GluN1 and two GluN2B ATD connected with specific 36 aa long linker peptides^25^ (Fig. 1B). The latter has the potential to assemble as a dimer of GluN1-GluN2B heterodimers as observed in the crystal structure of the entire ATD of NMDARs containing GluN1 and GluN2B (Fig. 1A).^26,27^ Both constructs containing either a mouse or a human Fc part were well soluble (up to at least 0.7 mg/ml ≈ 4 μM) and had the expected size of about 150 kDa (Fig. 1C).

**Fig. 1.**
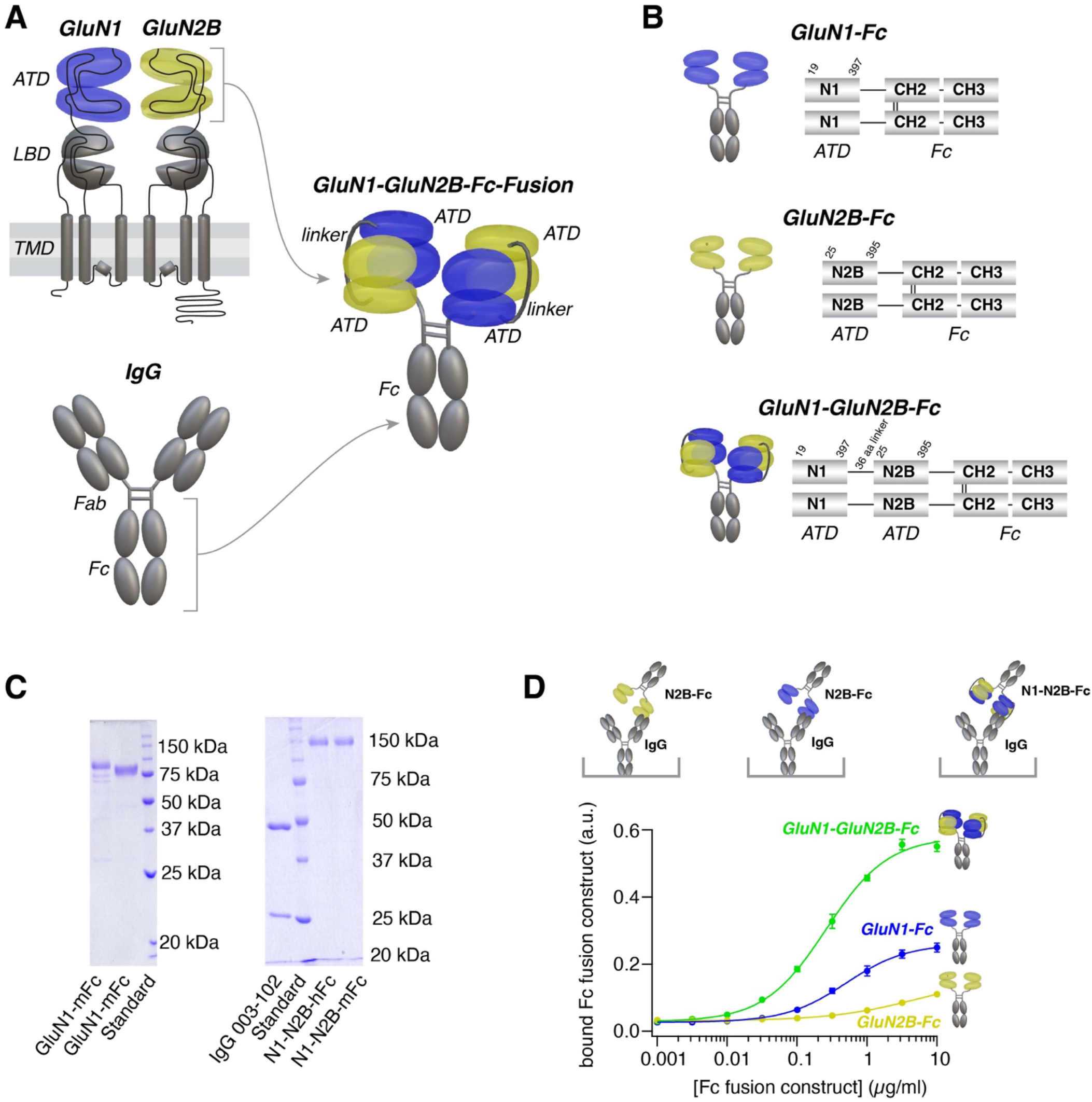
An engineered ATD-Fc-fusion construct has high affinity for pathogenic anti- NMDAR antibodies. (A) Illustration of the GluN1 and GluN2B subunits of the NMDAR (modified from^27^), an IgG molecule and a fusion construct containing two ATDs of GluN1, two ATDs of GluN2B, and an IgG Fc part. (B) Illustration of the three Fc-fusion constructs and their domains. (C) Sodium dodecyl sulfate polyacrylamide gel electrophoresis (SDS-PAGE) with Coomassie staining of fusion constructs (mFc: mouse Fc; hFc: human Fc). (D) *Top:* Illustration of the fusion constructs GluN1-Fc (N1-Fc), GlunN2B-Fc (N2B-Fc), and GluN1-GlunN2B-Fc (N1-N2B-Fc) bound to IgG, which was coated to an ELISA micro titer plate. *Bottom:* The amount of fusion constructs bound to monoclonal human IgG (003-102) plotted against the concentration of the fusion constructs containing the mouse Fc. The amount of the bound fusion constructs was detected with HRP conjugated secondary antibody against mouse Fc.

To test the binding capability of the constructs to pathogenic antibodies, we investigated their binding to a high-affinity monoclonal anti-NMDAR IgG isolated from a patient with NMDAR encephalitis (clone #003-102; ref.^28^). Coating the monoclonal anti-NMDAR IgG on a microtiter plate for an ELISA assay, titrating the concentration of the constructs, and subsequent immunostaining for bound constructs revealed the best binding interaction for the GluN1- GluN2B-Fc fusion construct (Fig. 1D). Similar results were obtained with a construct in which the order of the ATDs was changed (GluN2B-GluN1-Fc; data not shown). We selected the GluN1-GluN2B-Fc fusion construct as our leading molecule for further studies. Binding affinity of the GluN1-GluN2B-Fc fusion construct to the monoclonal anti-NMDAR IgG (003- 102; ref.^28^) was measured with biolayer interferometry (BLI, see Materials and methods). The measured dissociation constant (K_D_) was 29 nM, which is similar to the previously determined affinity of the binding of monoclonal anti-NMDAR IgG (003-102) to NMDARs (c_50_ = 1.2 μg/ml = 8 nM).^29^ These data indicate that a pathogenic anti-NMDAR autoantibody binds to the engineered GluN1-GluN2B-Fc fusion construct with high affinity.

### The ATD-Fc-fusion construct prevents binding of mono- and polyclonal pathogenic antibodies to NMDARs

To test the ability of the GluN1-GluN2B-Fc fusion construct to neutralize pathogenic antibodies, we tested its ability to prevent binding of pathogenic antibodies to NMDARs. As a first simple and robust approach, we used the recombinant ATDs of the GluN1-GluN2B-Fc fusion construct itself to serve as a NMDAR surrogate and coated it to a microtiter plate for an ELISA assay (Fig. 2A). A solution with a mixture of the pathogenic antibody with a constant concentration and varying concentrations of the fusion construct was added to the ELISA plate. After incubation, the plates were washed and the autoantibody binding to the immobilized fusion constructs was quantified. Adding increasing concentrations of the fusion construct prevented binding of the monoclonal anti-NMDAR IgG (003-102; ref.^28^) to the immobilized ATDs of the fusion constructs (Fig. 2B). The complete neutralization of the pathogenic antibodies suggests that two fusion constructs can bind to an antibody (one per Fab part) without steric hindrance.

**Fig. 2.**
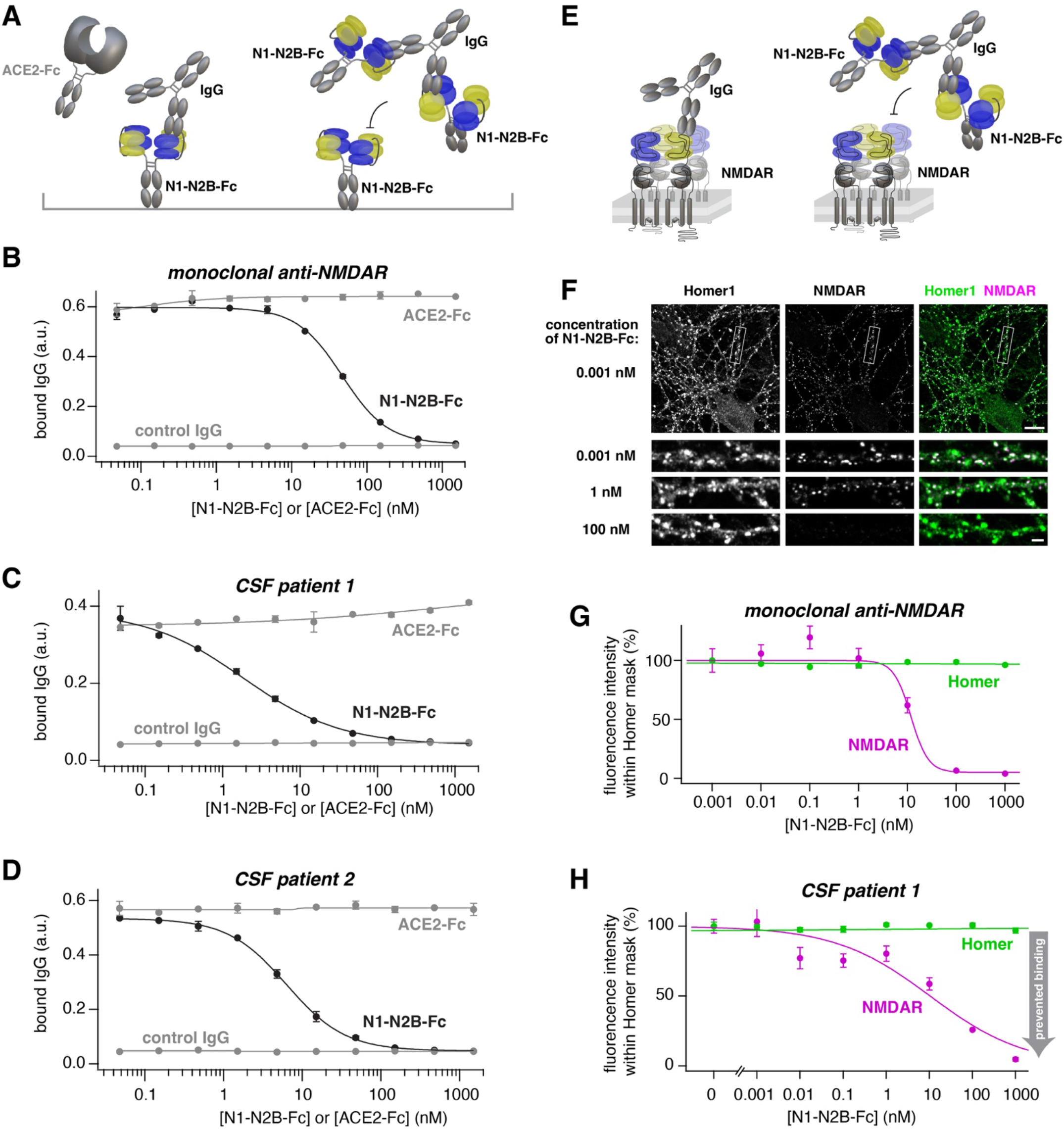
The ATD-Fc-fusion construct prevents binding of mono- and polyclonal pathogenic antibodies to NMDARs. (A) Illustration of GluN1-GlunN2B-Fc (N1-N2B-Fc) fusion construct coated to an ELISA micro titer plate with ACE2-Fc in solution and bound IgG (*left*). In addition, an IgG neutralized by two N1-N2B-Fc fusion constructs is shown (*right*). (B) IgG (monoclonal 003-102) bound to the coated GluN1-GlunN2B-Fc (N1-N2B-Fc) fusion constructs plotted against the concentration of the GluN1-GlunN2B-Fc (N1-N2B-Fc) fusion construct in solutions (*black*) or plotted against the concentration of a control fusion construct in solutions (ACE2-Fc; *grey*). The solutions always contained 1.5 nM of the monoclonal IgG. Bound control IgG (Palivizumab) is also shown (*grey*). The amount of bound IgG was quantified by labeling with a secondary antibody (mean ± SEM of measured values per n = 2 wells of the titer plate is shown) (C) Same as B but using CSF from patient 1 and from a control patient. (D) Same as B but using CSF from patient 2 and from a control patient. (E) Illustration of a NMDAR in a double lipid membrane of a neuron with bound IgG (*left*) and an IgG bound to two N1-N2B-Fc fusion constructs (*right*). (F) Example confocal fluorescence images of neurons stained for NMDARs (*magenta*) and Homer1 (*green*) treated for 24 h with CSF from patient 1 (1:20) and 0.001, 1, or 100 nM of the GluN1-GlunN2B-Fc (N1-N2B-Fc) fusion construct. Scale bar 10 μm (*top row*) and 2 μm (*lowest three rows*). (G) Average fluorescence intensity within the Homer1 masks for stained NMDAR (*magenta*) and Homer1 (*green*) plotted against the concentration of the GluN1-GlunN2B-Fc (N1-N2B- Fc) fusion construct in the solutions (mean ± SEM of the median fluorescence intensity per n = 10 images is shown; for each image, the median of the average pixel fluorescence of ∼500 synapses was calculated). The solutions always contained 1 nM of the monoclonal anti-NMDAR IgG (003-102). (H) Same as G but using CSF from patient 1 (1:20) for binding and staining of NMDARs.

To investigate whether the construct also prevents binding of a broader repertoire of pathogenic IgGs to the ATD of the NMDAR, we repeated the analysis with clinically relevant cerebrospinal fluid (CSF) samples containing high titer of NMDAR autoantibodies of two different patients (‘patient 1’ and ‘patient 2’). Consistent with the prevented binding of the monoclonal IgG, the construct prevented binding of the patients’ pathogenic antibodies to the recombinant ATDs (Fig. 2C and D). The half-maximal inhibitory effect was obtained at a concentration of 2.6 and 6.1 nM of the GluN1-GluN2B-Fc fusion construct for the CSF from patient 1 and 2, respectively. The constructs prevented IgG binding specifically, because increasing concentrations of a control Fc fusion construct (ACE2-Fc, see Materials and methods) did not impact IgG binding (Fig 2B-D) nor did control antibodies (Fig 2B) or CSF from a control patient (Fig. 2C and D) bind to the constructs.

We next tested the ability of the GluN1-GluN2B-Fc fusion construct to inhibit binding of pathogenic antibodies to synaptic NMDAR of fixed dissociated rat hippocampal neurons (Fig. 2E). The postsynaptic density of excitatory synapses was marked with anti-Homer1 antibodies and the NMDARs were stained with a monoclonal anti-NMDAR antibody (003-102; ref.^28^; Fig. 2F). With increasing concentrations of the fusion construct, the binding of anti-NMDAR IgG to Homer1-positive synapses was prevented (Fig. 2G). The half-maximal inhibitory effect was obtained at an 11-fold higher concentration of the construct compared with the concentration of the antibody. Identical analyses with CSF from patient 1 demonstrated that the construct prevents binding of patients’ pathogenic antibodies to synaptic NMDARs (Fig. 2H). The half-maximal inhibitory effect was obtained at a concentration of 10.4 nM GluN1-GluN2B-Fc fusion construct. These data demonstrate that the GluN1-GluN2B-Fc fusion construct can inhibit binding of mono- and polyclonal pathogenic antibodies to recombinant ATDs of NMDARs as well as synaptic NMDARs in hippocampal neurons.

### The fusion construct prevents NMDAR internalization in rodent and human neurons

A hallmark of the pathophysiology of NMDAR encephalitis is the internalization of NMDARs due to binding of bivalent autoantibodies.^7-9^ We therefore investigated NMDAR internalization in dissociated hippocampal neurons of rats (Fig. 3A). CSF from patient was applied for 24 h to the culturing medium of the neurons. Subsequently, NMDARs on the cell surface were stained with immunohistochemistry (see Materials and methods). The NMDAR signal within Homer1- postive excitatory synapses was reduced in a dose-dependent manner. Neurons treated with CSF of patient 1 at a dilution of 1:20 and 1:100 reduce the NMDAR intensity to 22% and 43% compared to neurons treated with control CSF, respectively, indicating profound NMDAR internalization (red data in Fig. 3B). Application of patient CSF and 100 nM of the GluN1- GluN2B-Fc fusion construct prevented receptor internalization (green data in Fig. 3B; median 108% and 98% for 1:20 and 1:100 dilutions, respectively). The fusion construct had no significant effect on the receptor density when applied for 24 hours with control CSF (grey data in Fig. 3B).

**Fig. 3.**
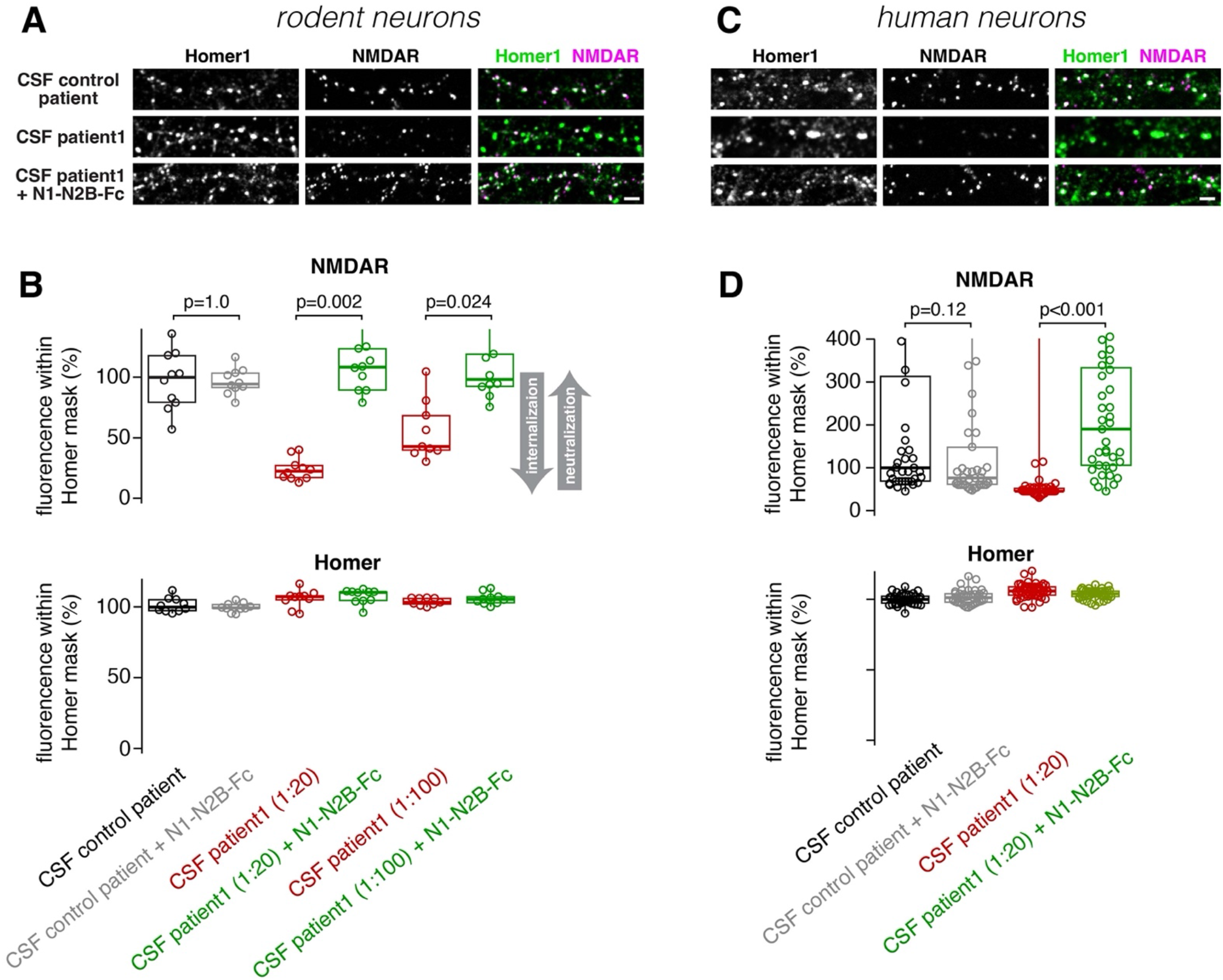
The fusion construct prevents NMDAR internalization in rodent and human neurons. (A) Example confocal fluorescence images of neurons stained for NMDARs (*magenta*) and Homer1 (*green*) after 24 h incubation with control CSF at a 1:20 dilution (*top row*), CSF from patient 1 at a 1:20 dilution (*middle row*), or CSF from patient 1 at a 1:20 dilution and 100 nM of the N1-N2B-Fc fusion construct (*lowest row*). Scale bar 2 μm. (B) Each data point represents the median of the average fluorescence pixel intensity within ∼500 Homer1-positive synapses per image for a staining against NMDARs (*top*) and Homer1 (*bottom*) of neurons treated for 24 h with control CSF at a 1:20 dilution (*black*), control CSF at a 1:20 dilution and 100 nM of the N1-N2B-Fc fusion construct (*grey*), CSF from patient 1 at a 1:20 dilution (*red*), CSF from patient 1 at a 1:20 dilution and 100 nM of the N1-N2B-Fc fusion construct (*green*), CSF from patient 1 at a 1:100 dilution (*red*), and CSF from patient 1 at a 1:100 dilution and 100 nM of the N1-N2B-Fc fusion construct (*green*). The data were normalized to the median value of neurons treated with control CSF. The indicated p values are from Dwass-Steel-Critchlow-Fligner pairwise comparisons (see Materials and methods). Note that some points are outside the plotted range. (C) Corresponding example images of human iPSC-derived neurons as shown in panel A. Scale bar 2 μm. (D) Corresponding analysis for human neurons as shown in panel B.

In addition, we repeated the NMDAR internalization assay in cultured neurogenin2-induced human iPSC-derived excitatory neurons (Fig. 3C).^30^ Despite the low density of NMDAR receptors in these cultured human neurons,^30^ we resolved an NMDAR internalization upon 24 h of incubation with patient CSF, which was inhibited by the GluN1-GluN2B-Fc fusion construct (Fig. 3D). These data indicate that the fusion construct blocks receptor internalization triggered by autoantibodies of patients with NMDAR encephalitis in both rodent and human neurons.

### The fusion construct stabilizes synaptic NMDAR currents

To functionally test the neutralization of pathogenic antibodies, we measured pharmacologically isolated spontaneous postsynaptic NMDAR currents in disassociated rat hippocampal neurons (Fig. 4A and B and S1). In neurons treated with CSF from patient 1 for 24 h, the postsynaptic currents were dramatically reduced compared with application of control CSF (Fig. 4B and C). In contrast, application of patient CSF and the fusion construct did not cause a significant reduction in the amplitude of postsynaptic currents (Fig. 4B and C). We observed an altered network activity in the cultured neurons treated with patient CSF, which could impact postsynaptic current amplitudes due to presynaptic vesicle depletion.^31^ We therefore repeated the analysis and analyzed only the very largest current amplitude per cell and obtained similar results (Fig. 4D). These data demonstrate that the newly-developed GluN1- GluN2B-Fc fusion construct inhibited the pathological reduction of synaptic NMDAR currents triggered by autoantibodies of patients suffering from NMDAR encephalitis.

**Fig. 4.**
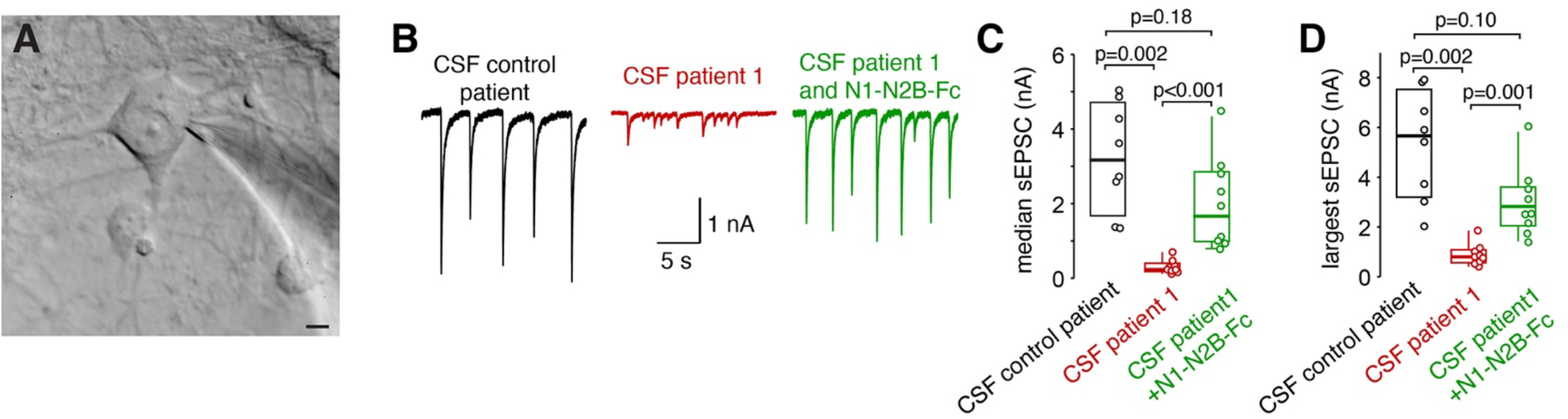
The fusion construct stabilizes synaptic NMDAR currents. (A) Example image of differential interference contrast (DIC) microscopy of a cultured hippocampal neuron (DIV 16) during whole-cell patch-clamp recording. Scale bar 5 μm. (B) Examples of pharmacologically isolated spontaneous NMDAR currents of neurons incubated for 24 h with control CSF (*black*), CSF from patient 1 at a 1:20 dilution (*red*), and CSF from patient 1 at a 1:20 dilution and 100 nM of the N1-N2B-Fc fusion construct (*green*). (C) Each data point represents the median amplitude per neuron of the spontaneous excitatory postsynaptic currents (sEPSCs). The indicated p values are from Dwass-Steel-Critchlow- Fligner pairwise comparisons (see Materials and methods). Note that some points are outside the plotted range. (D) Corresponding analysis as in panel C but each data point represents the very largest amplitude per neuron of the spontaneous excitatory postsynaptic currents (sEPSC).

## DISCUSSION

In this work, fusion constructs with ATDs of the NMDAR and IgG Fc parts were engineered. The construct containing the ATDs of both GluN1 and GluN2B subunits of the NMDAR bound pathogenic autoantibodies of patients with NMDAR encephalitis with high affinity. These constructs provide an exogenous target for the pathogenic antibodies prohibiting the binding of these antibodies to endogenous NMDAR receptors, thus serving as a ‘baitbody’ for pathogenic autoantibodies.

We here report a novel approach towards a rapid neutralization of antibodies in patients with NMDAR encephalitis which may also be combined with regular immunotherapy. In principle, such constructs could be used for direct and also repetitive intrathecal application in severely afflicted patients in the most active disease state during intensive care.^12^ However, further studies are required to investigate the optimal regime of intrathecal applications of the constructs in mouse models of NMDAR encephalitis. These studies are complicated by the variable severity of the established models after passive transfer using direct intrahippocampal injection or intrathecal application with osmotic pumps and due to their comparably fast time course of recovery from disease symptoms after stopping antibody delivery. The use of bispecific, so-called ‘brain-shuttle’ antibodies^32,33^ might be an interesting option for systemic application thus overcoming the blood-brain-barrier impermeability for macromolecules. Another therapeutic potential of the constructs could be a peripheral application for pregnant mothers with systemic anti-NMDAR autoantibodies, which might have transient or permanent pathogenic effects on CNS development of the fetus.^34,35^

The here-developed constructs could also add to the panel of diagnostic tools to reliably detect CSF or serum antibodies to the NMDAR. The current routine diagnostic relies on cell-based assays and visual inspection. Although widely used in clinical practice, these tests depend on the experience of the investigator. Additional and more specialized tests, e.g. brain section and live neuronal immunostaining can be necessary to clarify antibody detection in uncertain cases.^36-38^ A conformationally stable construct that can be applied easily in ELISA assays as demonstrated here might complement the routine cell-based assays and therefore contribute to robust, objective, and fast detection of anti-NMDAR antibodies.

Our data provide novel insights into the pathophysiology of NMDAR encephalitis. We found that the ATD of GluN1 alone but not of the GluN2B alone provided an epitope for pathogenic antibodies. The importance of the GluN1 subunit for antibody binding is consistent with existing evidence that the pathogenic antibodies bind only to the GluN1 subunit.^5,39^ Surprisingly, we found that the ATDs of both GluN1 and GluN2B were required for epitopes with the highest affinity. Our data thus indicate that either the high-affinity epitopes for pathogenic anti-NMDAR are comprised of both subunits, or that the ATD of GluN1 increases its affinity due to the interaction with GluN2B. In the first studies on NMDAR encephalitis identifying the GluN1 subunit as the target epitope for antibodies,^5,39^ the NMDAR subunits were expressed in HEK cells and the binding of the pathogenic antibodies was investigated in permeabilized cells with immunohistochemistry. Subtle alterations in the affinity of the epitope of GluN1 cannot be detected with these approaches. On the other hand, our construct contains only the ATD of GluN1 but not the entire GluN1 subunit. Therefore, the increase in affinity upon adding the ATD of Glun2B could be limited to ATDs isolated from the rest of the subunits. However, the simplest interpretation of our data is that the dimerization with GluN2B subunits is necessary for the GluN1 main immunogenic region to generate epitopes of highest affinity. This conclusion seems furthermore consistent with the difficulty of previous attempts to generate active immunization models. An active immunization model has only recently been established by using conformationally-stabilized and fully-assembled tetrameric GluN1/GluN2B NMDARs.^24^

In summary, we tested our here-developed fusion construct in a series of assays that are reflecting the critical steps of disease pathophysiology. Our data provide, to the best of our knowledge, the first therapeutic tool to efficiently and specifically neutralize pathogenic autoantibodies of patients suffering from NMDAR encephalitis that does not interfere with NMDAR function itself. The here-established therapeutic option of providing an antigen on an IgG Fc part (baitbody) could be used to neutralize antibodies against other antigens causing autoimmune encephalitis.^3^

## ACKNOWLEDGEMENTS

We thank Claudia Binder for expert technical assistance.

## MATERIALS AND METHODS

### NMDAR encephalitis patients and CSF sample collection

CSF samples from NMDAR encephalitis patients (NMDAR antibody titer in CSF 1:100) were collected from Jena University Hospital. All the voluntary donors were informed about the project and gave their written consent. The use of CSF samples was approved by the ethical committee of Jena University Hospital (approval # 2019-1415) and the Technische Universität Braunschweig (Ethik-Kommission der Fakultät 2 der TU Braunschweig, approval number FV- 2020-02).

### Fusion protein and IgG production

The three NMDAR fusion protein constructs were subcloned into pCSE2.7-mIgG2a-Fc-XP (N1-Fc and N2B-Fc) or pCSE2.8-mIgG2a-Fc-Xp (N1-N2B-Fc) using NcoI/NotI (New England Biolabs, Frankfurt, Germany) for mammalian production in Expi293F cells. The ACE2-Fc fusion protein, which served as a control fusion protein, was produced in Expi293F cells according to the methods and data published in ref.^40^. For the anti-NMDAR IgG (003-102; ref.^28^) production, the published amino acid sequences (WO 2017/029299 AI) were used and the V-Genes were ordered as GeneArt DNA fragments (Thermo Fisher Scientific, Schwerte, Germany) and were subcloned into IgG vectors human IgG1 format by subcloning of VH in the vector pCSEH1c (heavy chain) and VL in the vector pCSL3l/pCSL3k (light chain lambda/kappa)^41^ adapted for Golden Gate Assembly procedure with Esp3I restriction enzyme (New England Biolabs, Frankfurt, Germany). Expi293F cells were cultured at 37°C, 110 rpm and 5% CO_2_ in Gibco FreeStyle F17 expression media (Thermo Fisher Scientific) supplemented with 8 mM Glutamine and 0.1% Pluronic F68 (PAN Biotech). The Transfection cell density was between 1.5 - 2×10^6^ cells/ml and viability at about 90%. For formation of DNA:PEI complexes, 1 μg DNA/ml transfection volume and 5 μg of 40 kDa PEI (Polysciences) were first diluted separately in 5% transfection volume in supplemented F17 media. DNA (1:1 ratio of the vectors for IgG production) and PEI was then mixed and incubated ∼25 min at RT before addition to the cells. After 48 h the culture volume was doubled by feeding HyClone SFM4Transfx-293 media (GE Healthcare) supplemented with 8 mM Glutamine. Additionally, HyClone Boost 6 supplement (GE Healthcare) was added with 10% of the end volume. Harvesting was performed one week after transfection by 15 min centrifugation at 1500xg.

### Protein purification

Protein purification was performed as described in ref.^40^ using 1 ml column on Äkta go (Cytiva), Äkta Pure (Cytiva), or Profinia System (BIO-RAD). All of the purifications were carried out according to the manufacturer’s manual.

### SDS-PAGE and Coomassie staining

All produced fusion constructs and the anti-NMDAR IgG (003-102) were analyzed by sodium dodecyl sulfate-polyacrylamide gel electrophoresis (SDS-PAGE) with Coomassie staining. A 12% sodium dodecyl sulfate-polyacrylamide gel was prepared according to manufacturer’s protocol (BIO-RAD). For the analysis 32 μl of the production supernatant was mixed with 8 μl 5xLaemmli buffer containing ß-Mercaptoethanol and incubated for 10 min at 95°C. After a short cooldown and 20 sec centrifugation, 15 μl of each sample was loaded onto the 12% gel. Additionally, 8 μl of the BIO-RAD Precision Plus Protein All Blue Standard marker was loaded. The gel was running at 25 mA and 300 V in SDS running buffer (25 mM Tris; 192 mM Glycin; 00.1% SDS). After the separation run, the gel was stained with Coomassie Brilliant Blue (0.05% Coomassie-Brilliant Blue R250; 10% acetic acid; 25% isopropyl) overnight. The gel was destained for 2 h on following day with 10% acetic acid.

### Binding ELISA

Binding ELISA on the monoclonal anti-NMDAR IgG (003-102; Fig. 1D) was performed by immobilizing 200 ng of the 003-102 IgG on 96 well microtiter plates (High binding, Costar) in phosphate-buffered saline (PBS) with a pH of 7.4 overnight at 4°C. Afterwards, immobilized wells were blocked with 350 μl 2% M-PBST (2% milk powder in PBS with 0.05% Tween20). After blocking, wells were washed three times using H_2_O 0.05% Tween20. Respective fusion constructs were then titrated 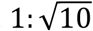 starting with 10 μg/ml. After 1 h incubation and washing three times with H_2_O 0.05% Tween20, binding was detected by using goat-anti-hIgG(Fc)-HRP (1:70000, A0170, Sigma). Binding was visualized by using TMB substrate: 20 parts TMB-A (30 mM potassium citrate; 1% (w/v) citric acid (pH 4.1) and 1 part TMB-B (10 mM TMB; 10% (v/v) acetone; 90% (v/v) ethanol; 80 mM H_2_O_2_). TMB reaction took place for 30 min and absorbance was measured at 450 nm with a 620 nm reference in an ELISA plate reader (Epoch, BioTek).

### Affinity measurement by Bio-Layer Interferometry

The affinity was measured by Bio-Layer Interferometry (BLI) using the Octet qKe (Fortebio/Sartorius GmbH, Göttingen, Germany). For this assay, anti-Human Fc-Capture (AHC) sensors were activated for 10 min in PBS. After that, the sensors were equilibrated in assay buffer (PBS containing 1% BSA and 0.05% Tween 20) for 60 s before anti-NMDAR IgG (003-102) was loaded onto the sensors at 10 μg/ml for 180 s. After a stable baseline measurement was established (60 s), loaded sensors were transferred to a 7-point N1-N2B-Fc dilution series (500, 158.1, 50, 15.8, 5, 1.58 and 0.5 nM). Association of the N1-N2B-Fc to the anti-NMDAR IgG was measured for 300 s. After that, the sensors were transferred into assay buffer (PBS) were the dissociation was measured for 600 s. Significant binding of the construct to an unloaded sensor was not detected. For data analysis, the reference measurement (0 nM) was subtracted from the other measurements and data traces ranging from 500 to 0.5 nM were used for modelling of the kinetic data using a 1:1 binding model (Octet qKe Data Analysis HT 11.0).

### Inhibition ELISA

The inhibition ELISA (Fig. 2A-D) was performed by immobilizing 200 ng N1-N2B-Fc on 96 well microtiter plates (High binding, Costar) and blocking with 2% milk powder. In parallel, on a separate blocked 96 well microtiter plates (Low binding, Costar) the N1-N2B-Fc construct was titrated 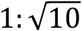 starting with a concentration of 1500 nM and going down to 0.005 nM. To the titrated fusion protein either 1.5 nM of monoclonal anti-NMDAR IgG (003-102) or 1:20 diluted CSF from NMDAR encephalitis patients was added and incubated for 1 h at RT. After incubation the N1-N2B-Fc autoantibody mixture was transferred to the immobilized N1-N2B- Fc and incubated for 1 h at RT. Afterwards, wells were washed three times using H_2_O 0.05% Tween20 and binding of NMDAR antibodies to the immobilized N1-N2B-Fc was detected by using goat-anti-hIgG(Fc)-HRP (1:70000, A0170, Sigma). Binding was visualized by using TMB substrate: 20 parts TMB-A (30 mM potassium citrate; 1% (w/v) citric acid (pH 4.1) and 1 part TMB-B (10 mM TMB; 10% (v/v) acetone; 90% (v/v) ethanol; 80 mM H_2_O_2_). TMB reaction took place for 30 min and absorbance was measured at 450 nm with a 620 nm reference in an ELISA plate reader (Epoch, BioTek).

### Laboratory animals

Dissociated hippocampal neurons were prepared using SPRD rats of either sex at postnatal (P) day 0 to 1. Cortical astrocytes were prepared using C57BL/6N mice of either sex at postnatal day 1∼3. Animals were kept in individually ventilated cages and received water and food ad libitum. All animal procedures were in accordance with the European (EU Directive 2010/63/EU, Annex IV for animal experiments), national and Leipzig University guidelines. All animal procedures were approved in advance by the federal Saxonian Animal Welfare Committee (T01/21 for mice, T29/19 for rat). In order to reduce the number of animals we tried to use for astrocyte cultures the remaining cells from other culture preparations in our laboratory whenever possible.

### Rat hippocampal cultures

Cultures from P0 to P1 rat were prepared according to ref.^42^ but with different coating of the coverslips, see below. Briefly, the hippocampus was dissected and enzymatically digested with Papain (Sigma) in the presence of DNAse (Sigma), followed by mechanical dissociation and centrifugation through a cushion of 4% bovine serum albumin (Sigma). These steps were completed using Hibernate medium (ThermoFisher). Cells were then plated onto Poly-D-Lysine (Sigma) coated coverslips in 24 well plates. For each coverslip 15,000 cells were allowed to settle in a 40 μl drop for about 30 min and then each well was filled with 500 μl growth medium: NeurobasalA/B27 (Invitrogen) supplemented with GlutaMax (0.25%, Invitrogen), Glutamine (0.25-0.5 mM, Sigma), Penicillin/Streptomycin (1:100, ThermoFisher) and heat-inactivated fetal calf serum (10%, Sigma). At day 2, 100 μl growth medium with AraC was added to inhibit overgrowth of astrocytes (final concentration: 1 μM, Sigma). Medium was partially exchanged on day 3 (480 μl) and day 7 (100 μl) with fresh maintenance medium: BrainPhys (StemCell), B27 (2%, Invitrogen), GlutaMax (0.25%, Invitrogen), Penicillin/Streptomycin (1%, ThermoFisher). Cultures were maintained for up to 3 weeks without any further medium change at 37°C and 5% CO2.

### Human induced neuron culture

The human induced pluripotent stem cell (iPSC) line BIHi005-A-24 (hPSCreg; https://hpscreg.eu/cell-line/BIHi005-A-24) was obtained from Berlin Institute of Health (BIH), Germany. HIV, HBV, HCV and mycoplasma were not detected by PCR. This cell line allows the induction of Ngn2 expression and the puromycin resistance gene by the addition of Doxycycline for differentiation into neurons.^30^ Human iPSCs were maintained in feeder-free culture using Matrigel (Sigma) coated plastic dishes and StemFlex medium (StemCell) without antibiotics. Frozen stocks were prepared in Bambanker HRM cryopreservation media (StemCell) and stored over liquid nitrogen. Upon reaching 50∼70% confluence iPSCs were harvested with Accutase (StemCell) and plated onto Matrigel (Sigma) coated 6 well plates at 200,000 cells/well in 2 ml/well StemFlex supplemented with ROCK inhibitor Y-27632 (5 mM, PeproTech) to inhibit apoptosis. The following three days (day 0-2) medium was replaced daily with 2 ml/well fresh differentiation medium: DMEM/F12/N2/NEAA (Invitrogen), penicillin/streptomycin (1%, Invitrogen), BDNF (10 ng/ml, PeproTech), 10 ng/ml NT-3 (10 ng/ml, PeproTech), mouse laminin (0.2 μg/ml, Invitrogen) and Doxycycline (2 μg/ml, Sigma). Puromycin selection was started by adding Puromycin (5 μg/ml, Sigma) on day 2. The iPSCs rapidly changed morphology towards a bipolar shape with elongated neurites. Because mouse astrocytes efficiently stimulate synaptogenesis in human induced neurons,^30^ cultures from P1 to P3 mice were prepared two to three weeks earlier with Trypsin digestion followed by mechanical dissociation and maintained in astrocyte medium: DMEM with 4.5 g/l glucose supplemented with pyruvate (1 mM, Sigma), heat inactivated fetal calf serum (10%, Sigma), penicillin/streptomycin (1%, ThermoFisher). They were replated two times to remove any mouse neurons. On day 3, neuron cultures were washed with HBSS to remove Puromycin and harvested with Accutase (StemCell). At the same time astrocyte cultures were harvested with Trypsin/EDTA (ThermoFisher). Cocultures were then plated onto Poly-D-Lysine (Sigma) and Matrigel (Sigma) coated coverslips in 24 well plates. For each coverslip 60,000 neurons and 12,000 astrocytes were allowed to settle in a 25 μl drop for about 30 min and then each well was filled with 500 μl neuron culture medium: NeurobasalA/B27 (Invitrogen), Penicillin/Streptomycin (1%, ThermoFisher), BDNF (10 ng/ml, PeproTech), 10 ng/ml NT-3 (10 ng/ml, PeproTech), mouse laminin (0.2 μg/ml, Invitrogen) and heat inactivated fetal calf serum (5%, Sigma). Half of the medium was replaced with neuron culture medium with Doxycycline on day 6, 10, and 14 and then twice a week. Doxycycline was continued until day 10. AraC (2 μM, Sigma) was added at day 12 for 48 h to inhibit overgrowth of astrocytes. During extended culture time an increase in osmolarity was observed. In this case, osmolarity of the medium, which was used for exchanging half of the medium, was diluted with H_2_O to obtain a final osmolarity of 300 mOsm in the medium. Cultures were maintained for up to 10 weeks at 37°C and 5% CO2.

### Immunohistochemistry on fixed neurons

To assess the inhibitory effect of the fusion construct N1-N2B-Fc on binding of CSF to NMDARs (Fig. 2E-H), the CSF of patient 1 (1:20 dilution) was pre-incubated with N1-N2B- Fc at various concentrations ranging from 1000 to 0.001 nM (in 1:10 dilution steps) at 4°C for 1 h. The cultured cells were fixated with 4% paraformaldehyde (PFA) in PBS at room temperature (RT) for 12 min, permeabilized with PBS and 0.3% Triton-X 100 at RT for 10 min and blocked in PBS, 5% normal goat serum (NGS), and 5% bovine serum albumin (BSA) for 1 h at RT. Afterwards, the pre-incubated CSF mixture was added to the cells and incubated at 4°C overnight. The bound anti-NMDAR antibodies were visualized by a corresponding fluorophore conjugated secondary antibody. To stain the postsynaptic density of excitatory synapses, neurons were incubated with a primary antibody (anti-Homer1) at 4°C overnight and subsequently with a corresponding fluorophore conjugated secondary antibody for 1 h at RT. Finally, all samples were stained with DAPI for 30 min at RT and embedded afterwards in Mowiol.

### NMDAR internalization

To measure NMDAR internalization (Fig. 3), the CSF of patient 1 (dilution 1:20 or 1:100) or control CSF was added to the culture medium of living rodent and human neurons for 24 h. In some experiments, 100 nM of the N1-N2B-Fc fusion construct was added simultaneously for 24 h. CSF and fusion constructs were not pre-mixed. To subsequently label NMDARs on the cell surface, a primary antibody against NMDAR receptors (003-102) was added to the medium for 30 min at 37°C, followed by a washing step with medium for 20 min at 37°C, another incubation with a corresponding fluorophore conjugated secondary antibody (CF568) for 30 min at 37°C, and another washing step with medium. Cells were then fixed with 4% PFA in PBS at room temperature for 12 min and permeabilized with PBS and 0.3% Triton-X 100 at RT for 10 min. A blocking step with PBS, 5% normal goat serum (NGS), and 5% bovine serum albumin (BSA) was performed prior to immunolabeling of the synaptic marker. The staining for Homer1 was performed as described above.

### Confocal imaging

All images were obtained using the confocal mode of an Expert Line Abberior microscope system with an Olympus (UPlanXApo) 60x oil objective with a numerical aperture of 1.42. The system contains 3 pulsed fluorescence excitation laser modules for wavelengths of 485, 561, and 640 nm, the corresponding filter cubes (GFP, Cy3, Cy5) and avalanche photodiode (APD) detectors for a highly sensitive signal detection. Each image was obtained by sequentially line scanning by switching between the needed excitation lasers line-wise during each recording. Each image contained a two channel-recording for stained Homer1 and NMDARs with 1252 × 1252 pixel size. Both channels were imaged with a dwell time of 6 μs/pixel and a twofold line accumulation. Based on the secondary antibody signals, the excitation laser power for each channel was adjusted to prevent over- or underexposure effects. Excitation laser power for Homer1 (secondary antibody, STAR Red, laser module 640 nm) was set to 0.3% and for NMDAR (secondary antibody, STAR Orange/CF568, laser module 561 nm) to 4%. The settings were kept constant throughout each set of experiments. Only the Homer1 signal was used for selecting a region for imaging within the cover slip.

### Image analysis

A custom-written macro of the software Fiji (ref.^43^) was used to generate a mask for Homer1- positive postsynaptic densities based on a prior adjusted threshold. The average pixel intensity of the NMDAR and Homer1 channel was quantified for each spot within the mask. For internalization experiments (Fig 3B and D), the median of the average pixel intensity of ∼500 Homer1-positive spots within an image was shown as an individual data point in the box plots. For the antibody binding curve (Fig 2G and H), corresponding median values per image were averaged and shown as the mean and the standard error of the mean (SEM).

### Electrophysiology

Rat hippocampal neurons [days in vitro (DIV) 15-17] were incubated with CSF from control patients, CSF from patient 1 at a 1:20 dilution or CSF from patient 1 at a 1:20 dilution and 100 nM of the N1-N2B-Fc fusion construct at 37°C for 24 h. Postsynaptic voltage-clamp recordings were performed at hippocampal neurons using a HEKA EPC10 amplifier (HEKA Elektronik, Lambrecht/Pfalz, Germany). Pipette solution for voltage clamp recordings contained (in mM): 130 CsMeSO_3_, 10 TEA-Cl, 10 HEPES, 5 EGTA, 3 Mg-ATP, 0.3 Na-GTP, 5 Na-Phosphocreatine, pH adjusted with KOH to 7.35 and osmolarity adjusted with sucrose to 300 mOsm. Series resistance (R_s_) was on average at 13.4 ± 0.6 MΩ and was compensated to remaining R_s_ of 4.4 ± 0.4 MΩ. Pipettes were pulled from borosilicate glass (Science Products, Hofheim, Germany) with a DMZ Universal Electrode (Zeitz Instruments, Martinsried, Germany) to resistances of 3 - 4 MΩ. Recordings of spontaneous NMDAR EPSCs were performed at a holding potential of -70 mV in magnesium-free extracellular solution containing (in mM): 145 NaCl, 2.5 KCl, 2 CaCl_2_, 10 HEPES, 15 glucose, pH adjusted by NaOH to 7.4. To isolate NMDA currents the extracellular solution was supplemented by 5 μM glycine, 10 μM NBQX, 10 μM SR95531, and 1 μM strychnine. A junction potential of 10 mV was not corrected. Recordings were performed at RT. To evaluate synaptic strength spontaneous events were measured for at least 2 minutes.

Current amplitudes were determined with the *waveform template matching* algorithm of the NeuroMatic plug-in^44^ (Version 3) for Igor Pro (WaveMetrics, Lake Oswego, OR, USA; Version 9). For each cell, either the median of all spontaneous EPSC amplitudes (Fig. 4C) or only the largest spontaneous EPSC amplitude (Fig. 4D) were shown as individual data points.

### Data analysis

The IC_50_ was calculated using a Hill equation

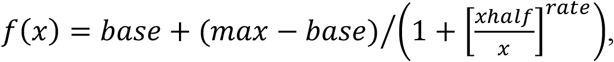

where *base* and *max* were constrained in some cases to 0 and 100% (for binding curves) or 100 and 0% (for inhibition curves), respectively. Fitting was done with Igor Pro (WaveMetrics, Lake Oswego, OR, USA; Version 9). For visualization of constructs (e.g. Fig. 1A), three-dimensional models were generated and rendered with blender (https://www.blender.org).

### Statistical analysis

For the data in Figs. 3B and D and 4B and C, non-parametric ANOVA (Kruskal-Wallis) tests revealed highly significant differences (p < 0.001). In the figures, the p values of non-parametric post-hoc tests (Dwass-Steel-Critchlow-Fligner pairwise comparisons) were provided. The calculations were performed with jamovi.^45^

### Data availability

Data that support the findings of this study are available from the corresponding author upon reasonable request.

### Resources table

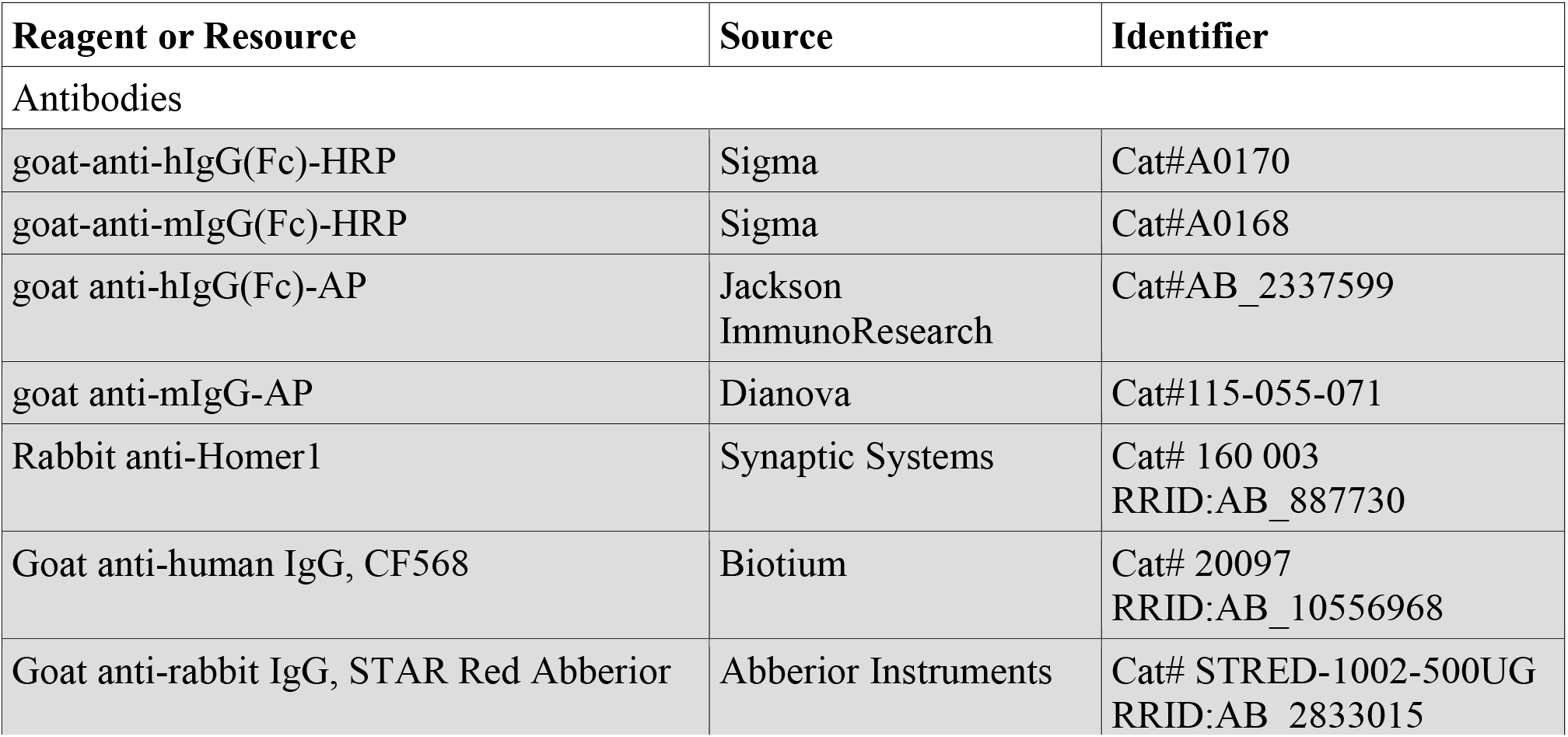
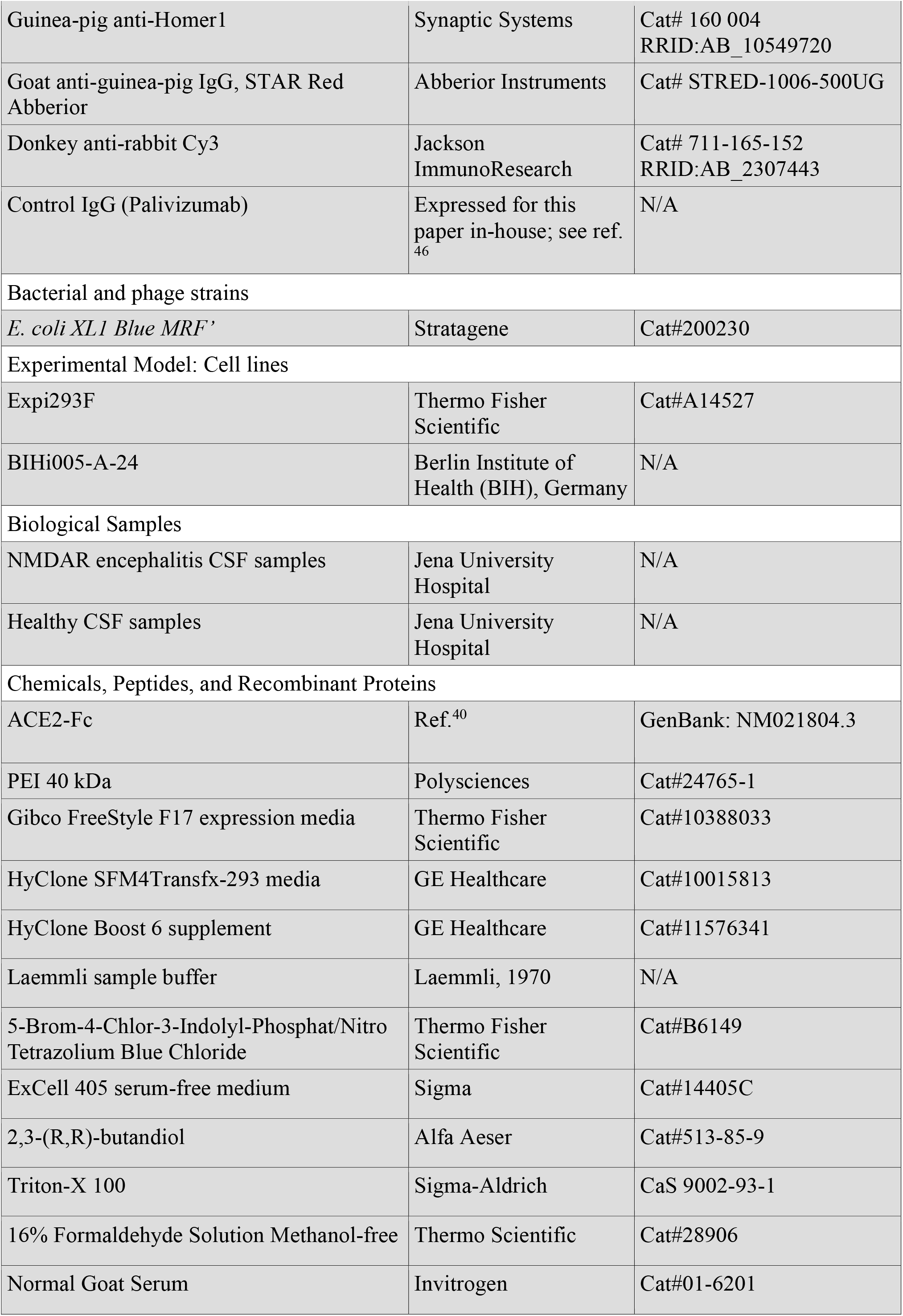
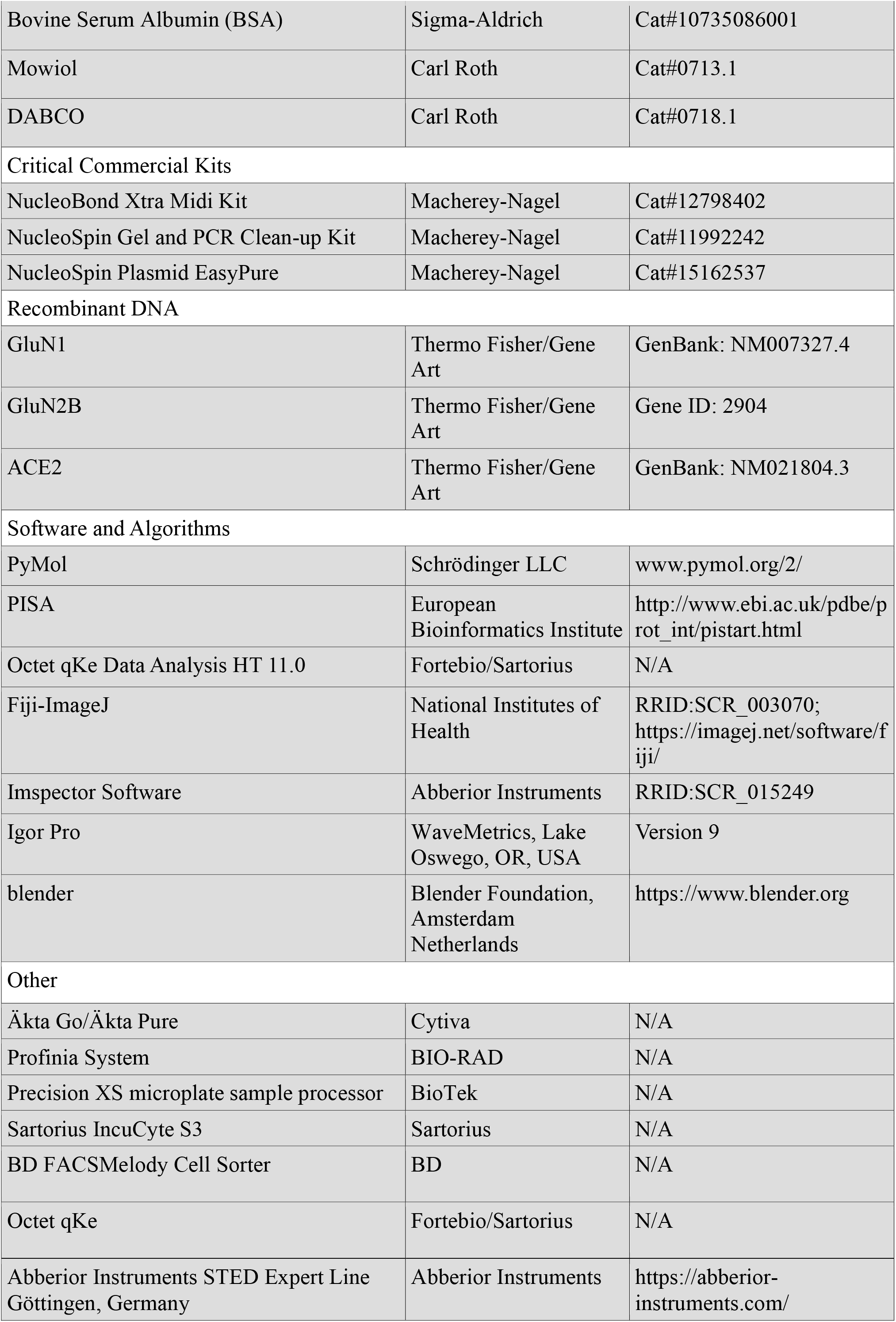

## FIGURES AND LEGENDS

**Fig. S1.**
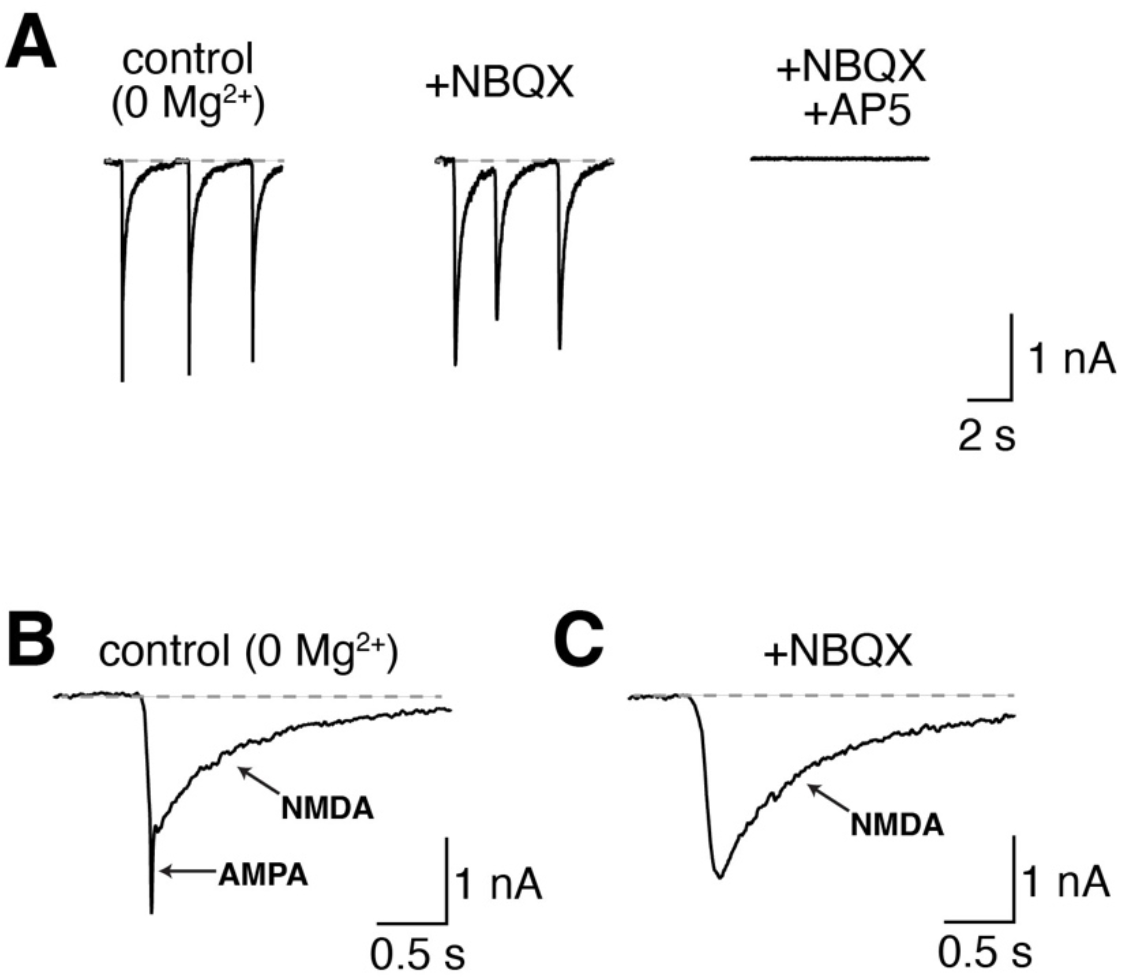
Pharmacological isolation of spontaneous NMDAR currents. (A) Examples of spontaneous excitatory postsynaptic currents (sEPSC) recorded at a holding potential of -70 mV in the magnesium-free extracellular solution (*left*), after addition of an AMPA receptor blocker (NBQX, *middle*), and after addition of a NMDAR blocker (AP5, *right*). (B) Example trace on a higher temporal resolution as shown in A. The AMPA receptor and NMDAR current components are indicated. (C) Example trace on a higher temporal resolution as shown in A after blocking the AMPA receptors with NBQX.

